## Supplementary Figures and Tables for "Collective dynamics support group drumming, reduce variability, and stabilize tempo drift"

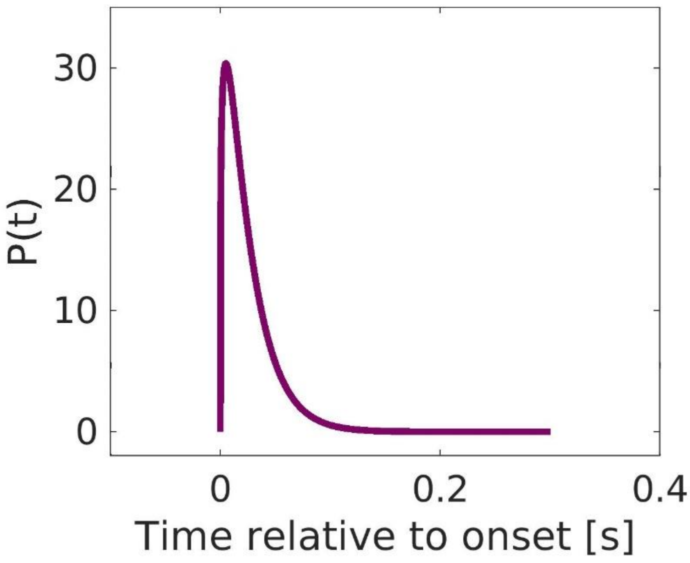


*Supplementary Figure S1*. The gamma distribution (*a*=1.25, *b*=.02) in the pulse-coupled model of group synchronization localizes the coupling in time, Eqs. 4-5.

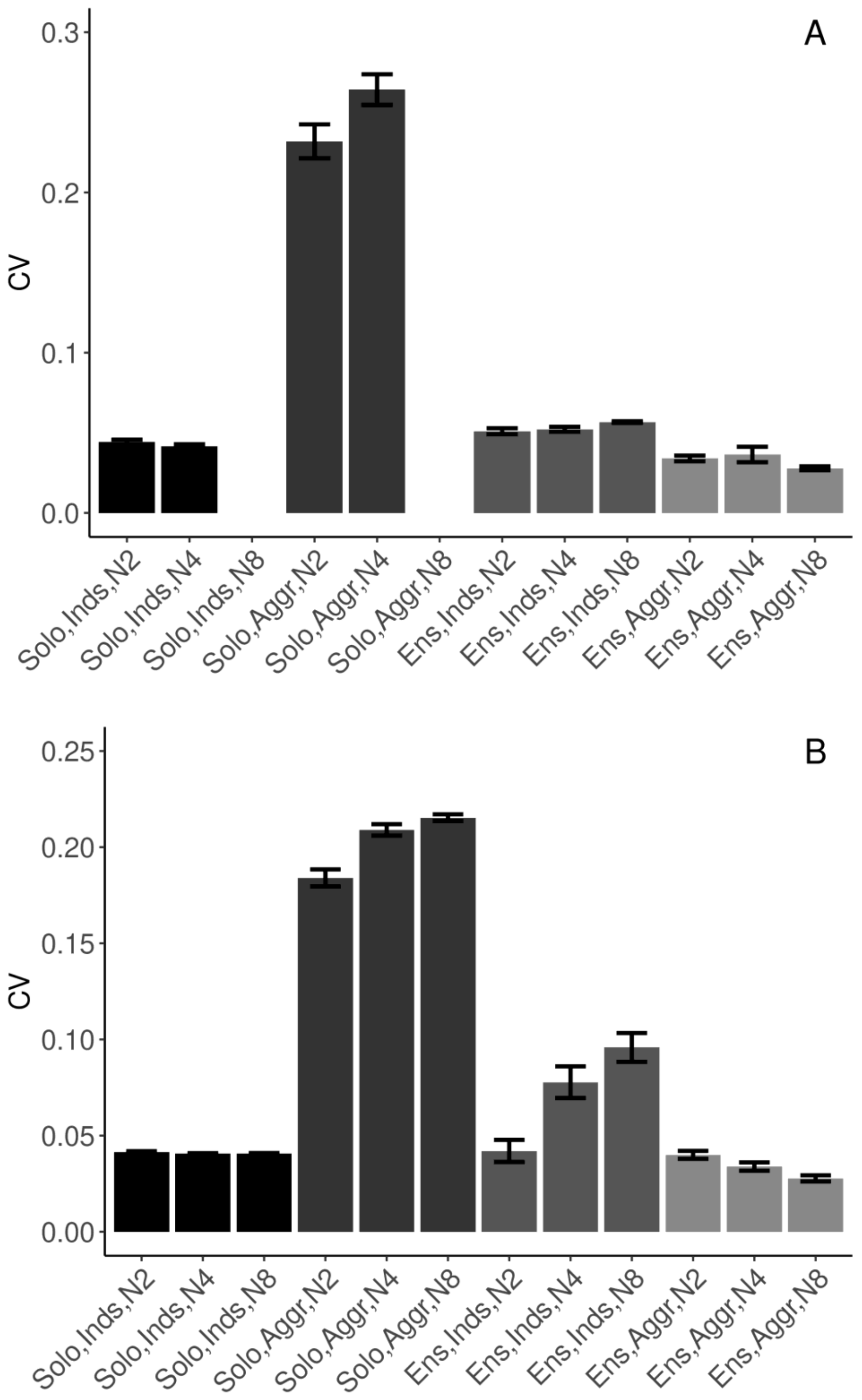


*Supplementary Figure S2*. Task performance measured in terms of mean (SE) coefficient of variation of IOIs. N2 = dyad; N4 = quartet; N8 = octet; Inds = individual participants; Agg = group-aggregate; Solo = solo condition; Ens = ensemble condition; no data collected in N8 Solo condition. (A) SCT drumming trials. (B) Simulations. As a sanity check, group-aggregate variability of IOIs in the solo (pseudo-group-ensemble) was much higher. A linear model was fitted to duet and quartet drumming trials (N2 and N4), the groups where the participants performed both in solo and ensemble condition. The statistical model confirmed that the group-aggregate variability in ensemble condition was lower than the pseudo-group-aggregate in solo playing condition [$\beta$=-.171, SE=.016, $t$=-10.56]. In addition, the pseudo-group-aggregate was higher than individuals' variability in solo conditions [$\beta$=.153, SE=.011, $t$=13.38]. There was an interaction between group size and the difference between individuals and group-aggregate. Pseudo-group-aggregate variability in solo trials increased as group size increased from duet to quartet [$\beta$=.017, SE=.004, $t$=4.67], that is, as more individual solo performances were put into calculating the mean field. In ensemble playing, group-aggregate variability decreased as group size increased from duet to quartet [$\beta$=-.016, SE=.005, $t$=-3.22], the more individuals playing together, the lower the variability of the group average.

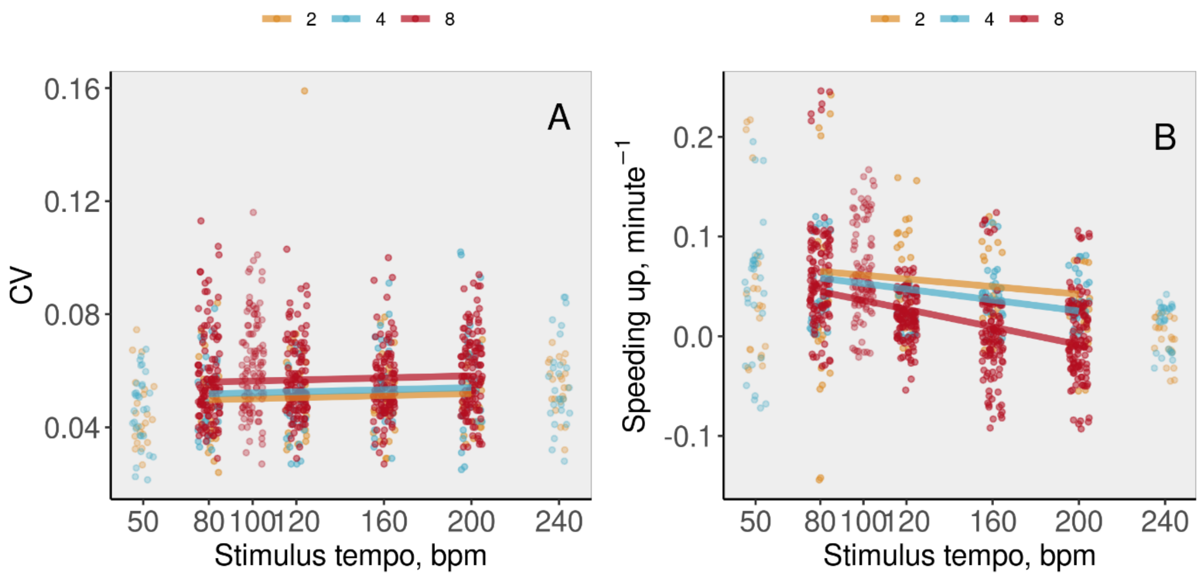


*Supplementary Figure S3*. Task performance in ensemble, SCT trials, measured in terms of A) variability and B) speeding up, defined as the linear slope of tempo over time. Individuals' data across group sizes (color-coded) and tempos are shown (abscissa jittered for visibility), along with the model fit. Note that tempos 50, 100, and 240 were not included in the statistical analysis and main text.

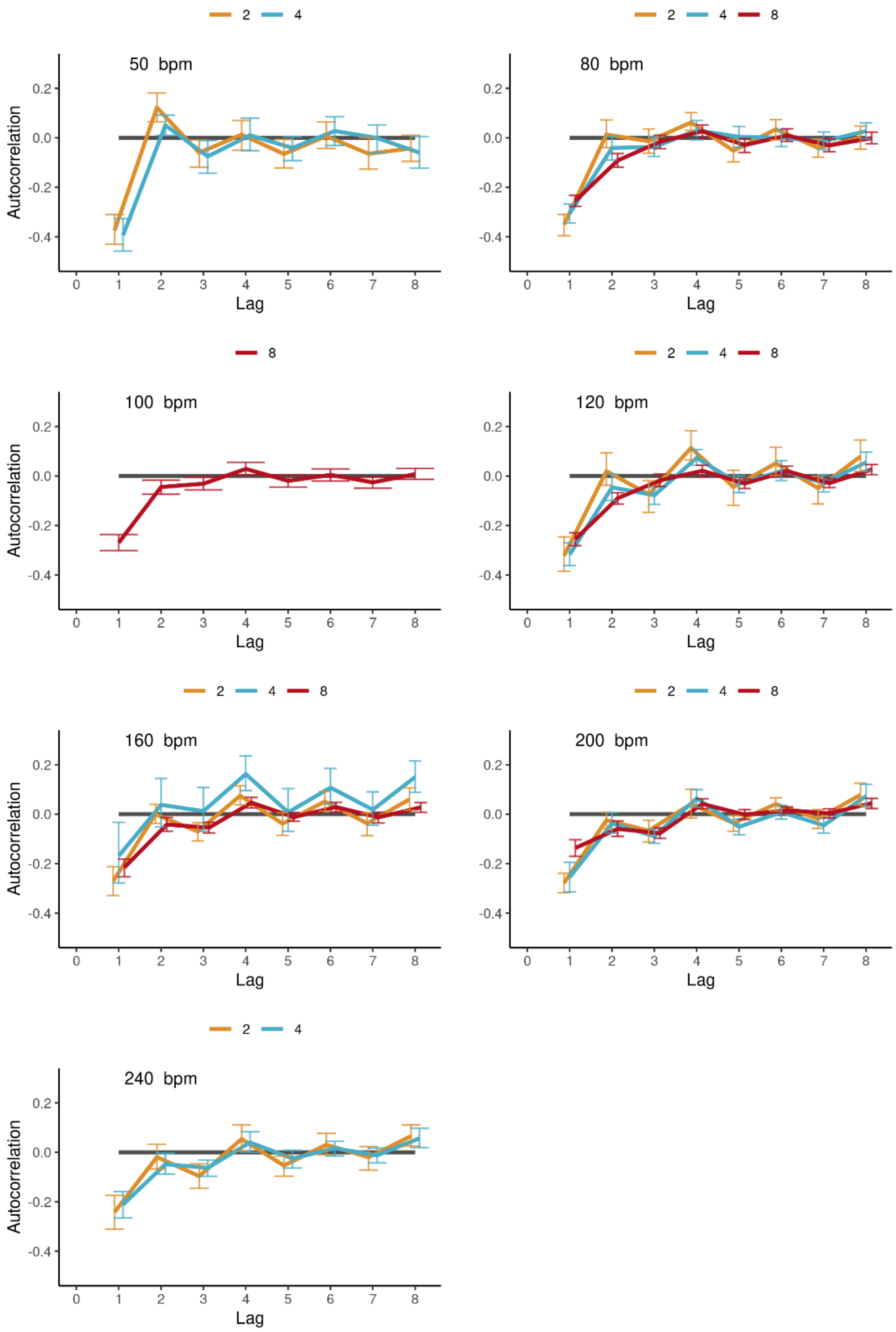


*Supplementary Figure S4.* Autocorrelations of individual IOIs in ensemble SCT drumming, averaged (SE) across individual participants' trials, shown separately per tempo (panels) and group size (color-coded lines). Tempos 50, 100, and 240 were not included in the statistical analysis in the main text.

Supplementary Table 1. Separate linear mixed-effects models were fitted for each of lags 1 to 8 of the auto-correlations of individuals (a) and group-aggregates (b) in the continuation phases of SCT trials, ensemble conditions. The predictors were *group size* ($N$) and *Tempo* (80, 120, 160, 200 bpm). For simplicity, the same full model was fitted in all cases for individuals, $Y_{ik}=(\beta_0+\sigma_{0g})+\beta_1 N_{ik}+\beta_2 Tempo_{ik}+\beta_3 N_{ik} Tempo_{ik}+\sigma_g$, where $i$ is participant, $k$ trial, and $g$ group. For the group-aggregate (b) where there was not enough data to fit models with all predictors, $Y_{ik}=(\beta_0+\sigma_{0g})+\beta_1 N_{ik}+\beta_2 Tempo_{ik}+\sigma_g$ was used. Significant coefficients ($p$<.05, Satterthwaite method) are in bold.

| Lag<br>Parameters | 1<br>*β/SE/t* | 2<br>*β/SE/t* | 3<br>*β/SE/t* | 4<br>*β/SE/t* | 5<br>*β/SE/t* | 6<br>*β/SE/t* | 7<br>*β/SE/t* | 8<br>*β/SE/t* |
|---|---|---|---|---|---|---|---|---|
| a) Individuals | | | | | | | | |
| $\beta_0$: Intercept | **-.425/.028/-15.19** | -.028/.023/-1.25 | *t*<1 | **.060/.019/3.14** | **-.05/.019/-2.65** | .028/.017/1.63 | **-.058/.018/-3.22** | *t*<1 |
| $\beta_1$: *N* | **.014/.005/3.01** | **-.012/.003/-4.46** | *t*<1 | **-.008/.002/-3.48** | *t*<1 | **-.004/.002/-1.99** | *t*<1 | **-.0066/.0029/-2.24** |
| $\beta_2$: Tempo | **.001/.0001/6.29** | .0002/.0001/1.64 | **-.0004/.0001/-3.83** | *t*<1 | *t*<1 | *t*<1 | **.0002/.0001/1.99** | **.0004/.0001/4.10** |
| b) Group-aggregate | | | | | | | | |
| $\beta_0$: Intercept | **-.228/.022/-10.34** | **.053/.016/3.34** | -.032/.020/-1.64 | **.11/.016/6.68** | **-.044/.015/-2.98** | **.033/.014/2.38** | *t*<1 | **.088/.015/5.81** |
| $\beta_1$: *N* | .0095/.0057/1.66 | -.005/.004/-1.18 | *t*<1 | **-.008/.004/-2.07** | .005/.004/1.20 | *t*<1 | *t*<1 | **-.009/.004/-2.38** |

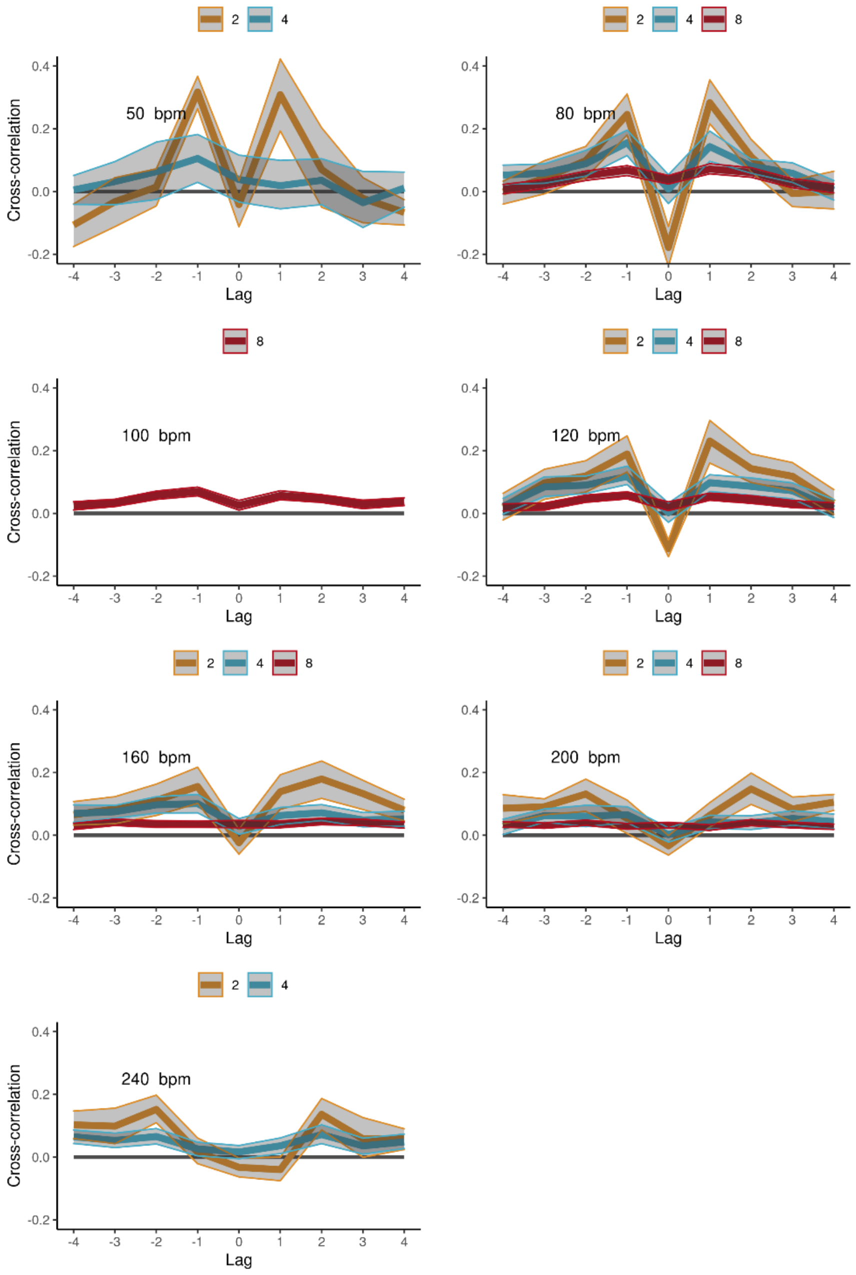


*Supplementary Figure S5*. Cross-correlations in the SCT task, ensemble conditions, averaged across participant pairs (+/- confidence intervals), per tempo (panels) and group size (color-coded lines). IOIs were aligned across participants and pre-whitened by filtering with an autoregressive model. Tempos 50, 100, and 240 were not included in the statistical analysis.

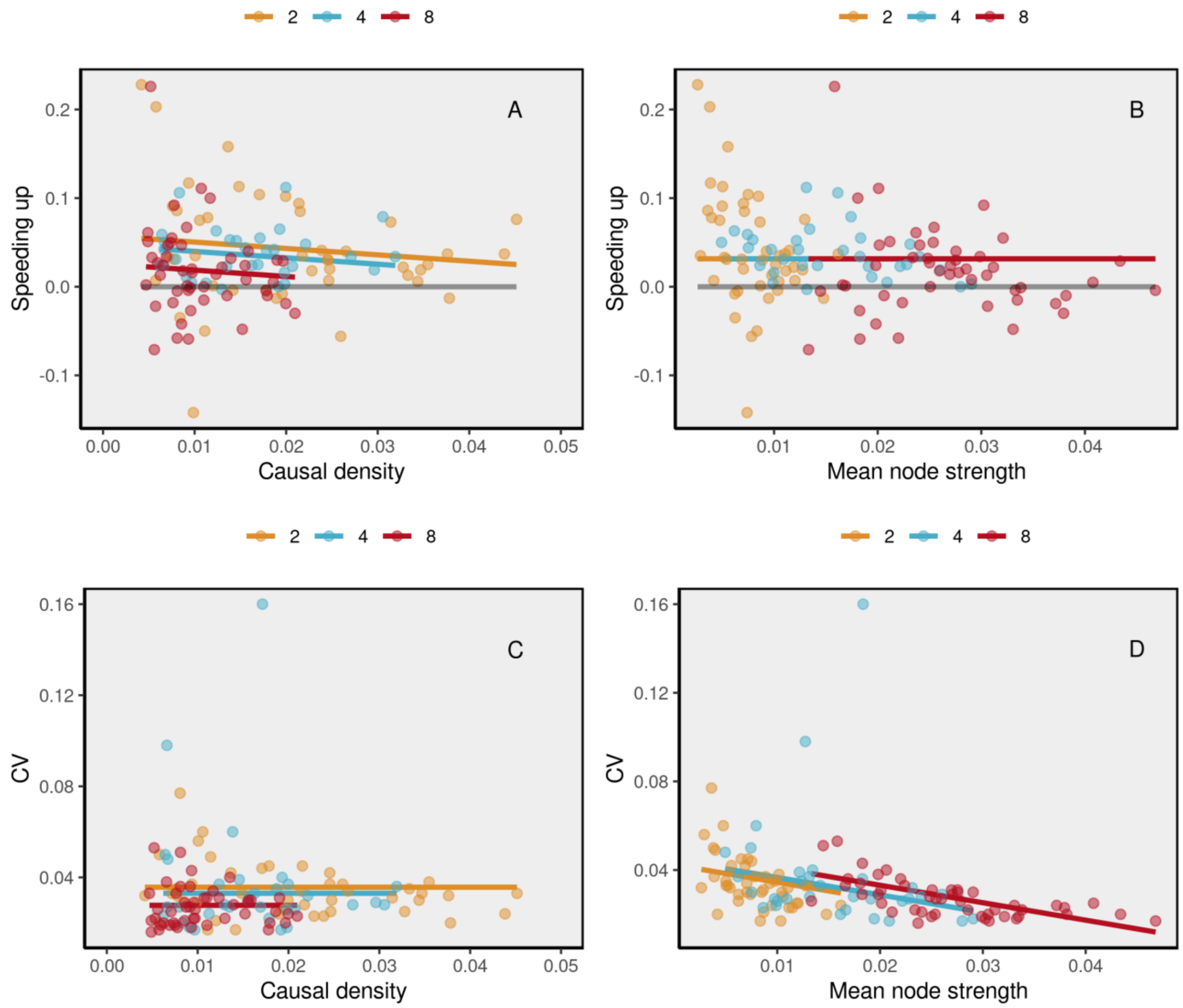


*Supplementary Figure S6*. Association between drumming performance measures and network properties in SCT ensemble trials.

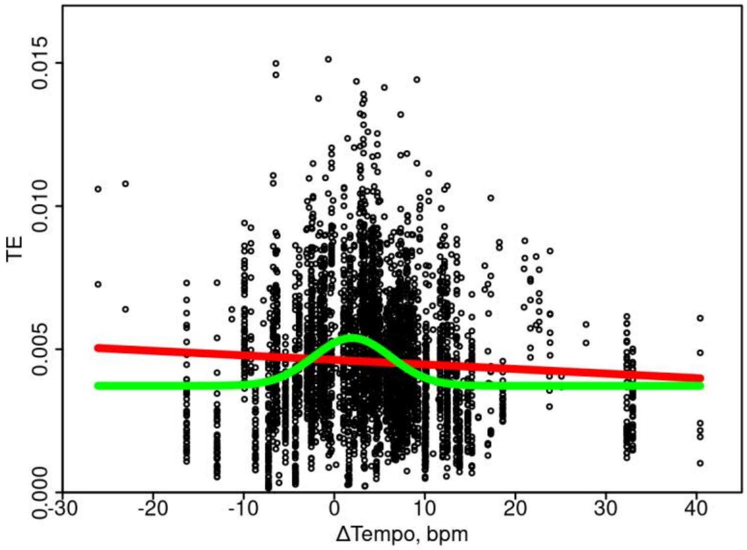


*Supplementary Figure S7.* TE is highest in SCT trials with minimal tempo increase relative to the instructed tempo. The Gaussian model (green) was selected over the linear (red) using model comparison (AIC and BIC), indicating that higher TE values tended to concentrate in the center of the distribution. Specifically, we compared a Gaussian and a linear model's ability to predict TE values from all pairs during continuation tapping (including all group sizes) from their respective changes in tempo (maximum likelihood estimation, mle in R). Indeed, the AIC method selected the Gaussian model (-39838.89) over the linear (-39566.81). The estimated 95% confidence intervals of model parameters indicated that the center of the Gaussian curve was just above zero, 1.56 to 2.37 bmp change in tempo, over the continuation phase (tempo changes in the data ranged from about -20 to 40 bmp), suggesting that best group coordination was achieved when participants were slowly increasing their tempo, or that slowly increasing the tempo allowed for best coordination.

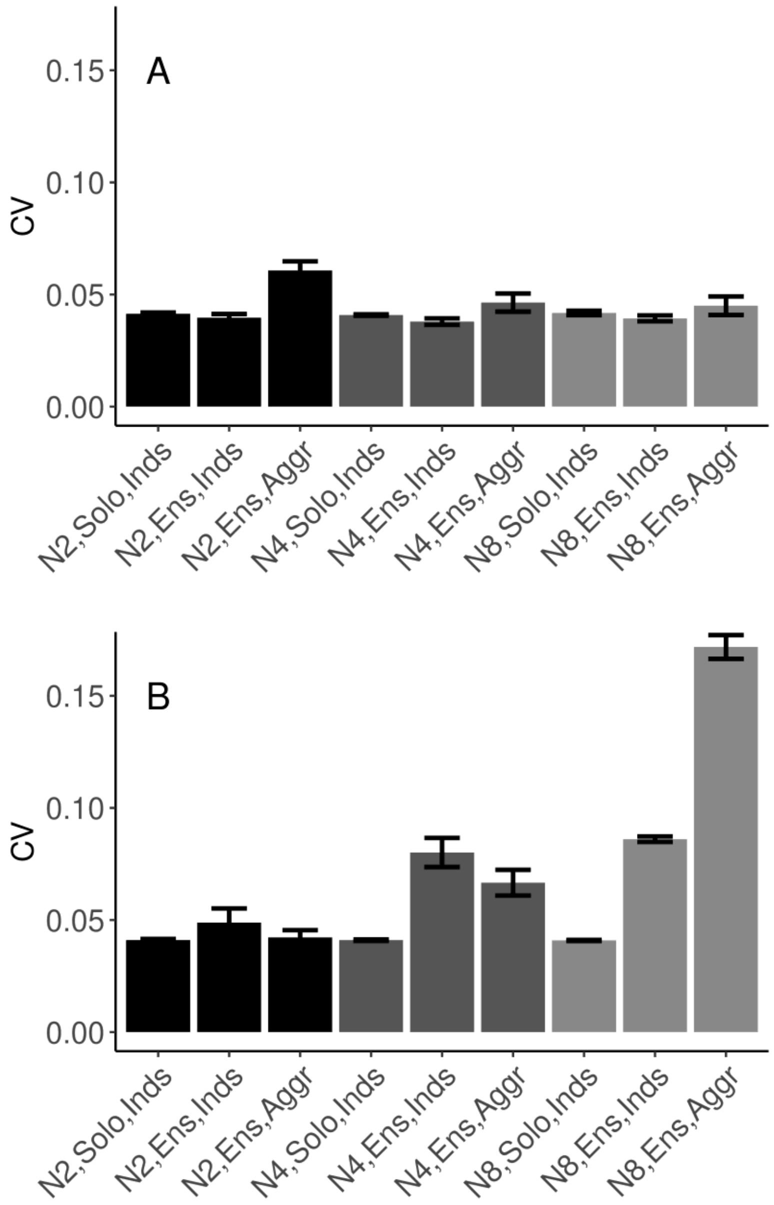


*Supplementary Figure S8*. Variability in alternative models of group synchronization. (A) The classical Kuramoto model with constant coupling term. (B) The pulse-coupled model, Eqs. 4-5, but with selective coupling in a ring topology, where units only interact with their immediate neighbors. In contrast to the pulse-coupled model developed in the present paper, these models do not exhibit the pattern of data across conditions found empirically, see Figures 4A-B.
